# How different AI models understand cells differently

**DOI:** 10.64898/2026.01.29.702682

**Authors:** Yubo Zhao, Duozhi Sun, Minsheng Hao, Yifan Xiong, Chen Li, Tieliang Gong, Lei Wei, Xuegong Zhang

**Author notes:** Correspondence to: Lei Wei < >, Xuegong Zhang < >.

## Abstract

AI single-cell foundation models (scFMs) are believed to be able to learn essential relations in cell transcriptomics with the attention modules in Transformer, but there is no method to reveal what they actually learned. We observed that different models may grasp different aspects of relations. To unravel the mystery, we propose scGeneLens, a framework for dissecting how scFMs perceive cells. We employed a sparse block attention to replace the original attention mechanism to concentrate attentions into a few dominant gene–gene relations, used attention propagation to trace how the relations propagate across Transformer layers, and used integrated gradients to disentangle the relative contributions of gene identity and expression in cell representations. We applied it to scFoundation and scGPT and show that they exhibit pluralistic perceptions of cells: scFoundation emphasizes relations among cell-type marker genes, resulting in stronger cell-type separability, whereas scGPT focuses more on genes involved in shared cellular pathways and core biological activities, leading to representations that generalize across conditions. The framework provides a unified lens for probing what scFMs learn about cells and offers actionable insights for the design of future cellular foundation models. Our code can be seen in https://anonymous.4open.science/r/scGeneLens-B771/.

## 1. Introduction

Transformer-based single-cell foundation models (scFMs), pretrained on large-scale scRNA-seq corpora, have emerged as powerful backbones for a wide range of downstream analyses (Bian et al., 2024). Representative models such as Geneformer (Theodoris et al., 2023), scBERT (Yang et al., 2022), scGPT (Cui et al., 2024), and scFoundation (Hao et al., 2024) share broadly similar Transformer-based architectures, but differ substantially in their pretraining objectives, input representations, and optimization strategies. As a result, architectural similarity does not imply that these models encode cell states in the same way.

Consistent with this distinction, recent benchmarks and zero-shot evaluations have shown that no single scFM consistently outperforms others across gene-level and cell-level tasks, with performance strongly depending on task definitions, evaluation protocols, and dataset characteristics (Wu et al., 2025; Kedzierska et al., 2025). These heterogeneous behaviors suggest that different scFMs rely on distinct inductive biases when organizing gene-level information into cell representations. However, most existing evaluations treat scFMs as black boxes and provide limited insight into how these differences arise internally.

A natural entry point for probing such differences is the self-attention mechanism, which governs how information from thousands of gene tokens is weighted and integrated. In principle, attention patterns could reveal which genes or gene groups are prioritized when encoding cell states. In practice, however, attention in scFMs is notoriously difficult to interpret: the input context is extremely long, attention weights are dense and highly distributed, and their influence becomes increasingly opaque across layers. Naïve inspection of attention maps therefore often yields noisy or inconclusive interpretations.

To address these challenges, we propose scGeneLens, a model-agnostic framework for dissecting how scFMs perceive cells through structured attention analysis. scGeneLens minimally modifies the attention mechanism to reorganize dense self-attention into a compact set of dominant gene–gene dependencies, traces how these dependencies propagate across Transformer layers, and disentangles the relative contributions of gene identity and expression magnitude to cell representations. Here, block-sparse attention provides a structured abstraction of the original attention patterns by aggregating interactions among genes with similar contextual roles, making dominant gene–gene dependencies more interpretable without altering the underlying representational capacity.

We apply scGeneLens to two representative scFMs, scFoundation and scGPT, and uncover systematic differences in the biological signals they emphasize. scFoundation preferentially organizes representations around cell-type marker genes, yielding strong cell-type separability and attention propagation anchored to identity-defining modules. In contrast, scGPT places greater weight on expression-driven signals associated with shared pathways and core cellular activities, resulting in representations that generalize across biological conditions. These contrasting behaviors provide a mechanistic explanation for the divergent performance profiles of scFMs observed across tasks.

Together, our results show that scFMs with similar architectures can encode fundamentally different notions of cell state. By offering a unified and interpretable lens on attention-based representations, scGeneLens enables systematic comparison of inductive biases across models and provides actionable insights for the analysis, evaluation, and future design of single-cell foundation models.

## 2. Related Works

### 2.1. Efforts on Interpretation of scFMs

While scFMs are increasingly adopted in analyses, their internal mechanisms remain poorly understood, limiting reliable biological interpretation and practical deployment. Existing interpretability efforts primarily focus on hidden states and embeddings, for example by using sparse autoencoders to extract compact and human-interpretable features from model activations (Pedrocchi et al., 2025; Claye et al., 2025; Schuster, 2025). While these approaches provide useful summaries of learned representations, they operate on the final or intermediate features produced by the model and thus offer limited insight into the underlying mechanisms by which gene-level information is processed.

In contrast, the self-attention mechanism directly governs how signals from thousands of gene tokens are weighted, selected, and integrated to form cell representations. Without explicitly examining attention, it remains difficult to disentangle whether differences in representations arise from the choice of genes emphasized, the structure of their interactions, or subsequent nonlinear transformations. As a result, the attention mechanism represents a more proximal and mechanistically informative target for interpreting how scFMs understand cell states.

### 2.2. Attention-Based Interpretability for Transformers

In natural language processing (NLP), attention has long been explored as a tool for interpretation. However, attention weights from individual layers are often noisy and do not reliably reflect true importance (Jain & Wallace, 2019). Attention rollout and attention flow methods address this by aggregating attention across layers to estimate effective token influence. Such approaches have been shown to recover intuitive dependencies, for example highlighting subject–verb relations, and to support inspection of masked-word predictions by tracing influential context tokens (Abnar & Zuidema, 2020). These methods provide a general framework for interpreting deep, multi-layer attention models, but face substantial challenges when applied to the single-cell setting, where attention operates over thousands of gene tokens and attention weights are correspondingly more diffuse and harder to attribute.

### 2.3. Block Sparse Attention

As sequence length increases, the *O*(*L*^2^) computational and memory costs of dense self-attention become prohibitive, motivating block-structured sparsification strategies in long-context Transformers. These approaches group tokens into blocks and restrict each query to attend to a small subset of blocks, reducing complexity while preserving dominant contextual interactions. Representative methods include Native Sparse Attention (NSA), which combines local windows, selected global tokens, and compressed block summaries (Yuan et al., 2025), and MoBA, which dynamically routes queries to relevant key–value blocks via a mixture-of-experts mechanism (Lu et al., 2025).

Although primarily developed for efficiency, block-sparse attention methods share a common principle of reorganizing dense attention into structured, salient interactions. This principle is particularly relevant to scFMs, where each cell is represented by thousands of gene tokens and attention patterns are correspondingly large and diffuse. In this work, we adopt block-structured sparsification not for acceleration, but to render attention patterns compact and analyzable for studying gene–gene dependencies in scFMs.

## 3. Methods

We propose SCGeneLens, an analysis framework for dissecting how gene-level information is integrated through attention mechanisms in scFMs. scGeneLens comprises three tightly coupled components: (i) **scGeneLens-Attention**, a gated block-sparse attention module that reorganizes dense self-attention into analyzable gene–gene dependencies; (ii) a cross-layer **gene influence propagation** scheme that traces how attention-mediated dependencies accumulate across depth; and (iii) **integrated gradients** applied to model inputs to disentangle the relative contributions of gene identity and expression magnitude. Together, these components enable a systematic and interpretable dissection of attention-based representations in scFMs (Fig. 1).

**Figure 1:**
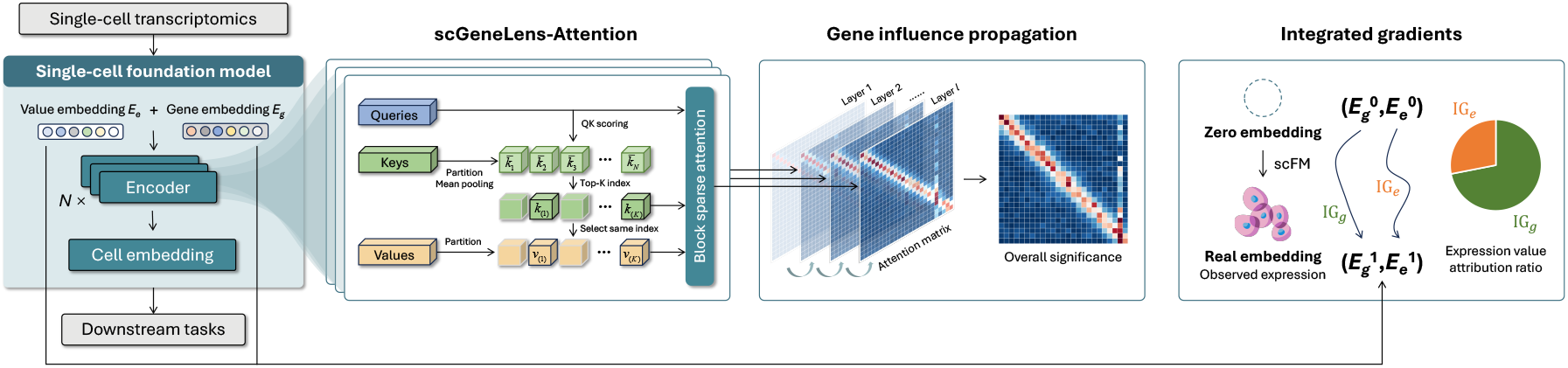
Overview of scGeneLens.

### 3.1. scGeneLens-Attention

To enable structured analysis of attention mechanisms in scFMs, inspired by block-sparse attention designs such as MoBA (Lu et al., 2025), we introduce **scGeneLens-Attention** as an architecture-preserving attention approximation that can replace dense global attention in scFM encoders for analysis.

Given an input gene representation **X** ∈ ℝ^*n*×*d*^, we follow the standard Transformer formulation and project it into query, key, and value tensors **Q, K**, and **V**. To enable structured attention analysis, gene tokens are partitioned into non-overlapping blocks: genes are ordered lexicographically with special tokens appended at the end, and the resulting sequence is split into consecutive blocks with a fixed chunk size *M* along the token dimension.

Accordingly, the key and value tensors are partitioned into *N* non-overlapping blocks:

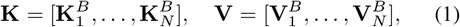

where each block 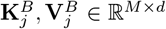 groups a contiguous subset of genes and serves as a coarse-grained unit for block-level attention analysis. When the input sequence length *n* is not divisible by *M*, the final block may contain fewer than *M* tokens.

For a query gene *g*_*i*_ with query vector **q**_*i*_, scGeneLens evaluates relevance at the block level rather than directly at the token level. Each block is summarized by its centroid:

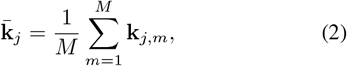

and a query–block relevance score is computed as

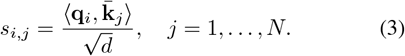

To induce sparsity, only the top-*k* scoring blocks are activated for each query:

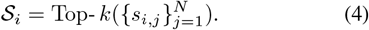

If *n* is not divisible by *M*, the final (short) block is always retained, as it contains special tokens placed at the end of the sequence that encode global information. This gating step confines attention to a limited set of relevant gene blocks, yielding a sparse and more analyzable attention pattern.

Token-level attention is then computed within the selected blocks. As gene expression profiles are treated as permutation-invariant sets, no causal mask is applied. Let 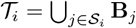 denote the union of tokens in the selected blocks. The final attention output for query **q**_*i*_ is given by

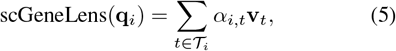

where *α*_*i,t*_ is obtained via a softmax over 𝒯_*i*_.

Together, scGeneLens-Attention reorganizes dense selfattention into a structured, block-sparse form that highlights dominant gene–gene dependencies while preserving attention computation within selected blocks.

### 3.2. Gene Influence Propagation

To analyze how scGeneLens-Attention propagates genelevel information across multiple encoder layers, inspired by attention rollout (Abnar & Zuidema, 2020), we introduce a **gene influence propagation** mechanism. This method traces how gene-level influences accumulate through depth by explicitly composing sparse attention transformations across layers.

At each layer *l*, attention is computed using *H* heads, yielding sparse attention matrices 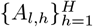. We first aggregate these matrices by mean pooling:

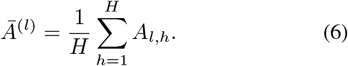

To account for residual connections that preserve a gene’s own contribution across layers, we add a scaled identity matrix to the aggregated attention. The resulting matrix is then row-normalized to ensure a consistent distribution of influence:

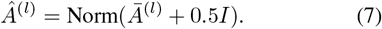

This step balances outgoing influence from each gene, enabling stable comparison of influence patterns across layers.

To capture cumulative gene influence, we recursively compose the balanced attention matrices across layers:

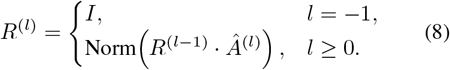

The resulting matrix *R* ^(*L*−1)^ represents the total integrated influence after *L* layers. In particular, 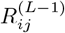 quantifies the relative contribution of the initial representation of gene *j* to the final refined representation of gene *i*.

Overall, gene influence propagation converts layer-wise sparse attention patterns into an interpretable influence map that captures how gene–gene dependencies derived by scGeneLens-Attention accumulate across depth.

### 3.3. Integrated Gradients

To disentangle the sources of information driving cell representations, we apply **integrated gradients** (Sundararajan et al., 2017) to quantify the relative contributions of gene identity and expression magnitude. In scFMs, each input token combines a gene embedding *E*_*g*_ and an expression value embedding *E*_*e*_, corresponding to gene identity and expression level, respectively, and integrated gradients attributes the model output to these two components. Since integrated gradients require a scalar output, we choose a scalar proxy objective *F* on the cell embedding *z*, specifically *F* (*z*) =∥ *z* ∥ _2_. Given a zero-embedding baseline 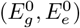 and a real embedding 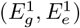, we compute attribution scores for gene identity and expression value as

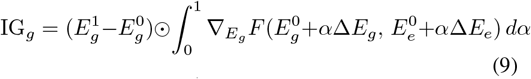

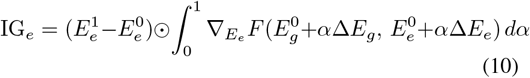

To summarize the model’s relative dependence on expression magnitude, we define an expression value attribution ratio

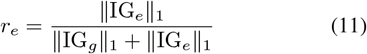

This analysis complements attention-based interpretation by distinguishing whether gene–gene dependencies and cross-layer influences are primarily driven by gene identity or expression magnitude, offering a finer-grained view of how scFMs encode cellular information. Model setup and parameters are described in Appendix A.1, and datasets are detailed in Appendix A.2.

## 4. Results

### 4.1. Differential Information Encoding in scFMs

#### 4.1.1. Intrinsically Preserved Cell Identity Information IN SCFoundation

We first evaluate scFoundation and scGPT in a cell-type annotation setting under supervised fine-tuning with scGeneLens. Both models are fine-tuned on a human pancreatic dataset (Segerstolpe et al., 2016) largely following their default and recommended fine-tuning protocols. For scGPT, we introduce a minor modification to the default setup by keeping the embedding layer frozen as our goal is to examine what information is already encoded by the pre-trained representations. We additionally evaluate three parameter-freezing strategies on scGPT with varying numbers of trainable parameters, including one aligned with the scFoundation update scheme (Table 1).

**Table 1:** Configurations and accuracies for cell-type fine-tuning. For scGeneLens, TopK = 6 and chunk size = 16. ✓: trainable; –: frozen; All: fine-tuning all layers; *L* − *k*: fine-tuning the *k*-to-last layer. EL: embedding layer; MLP: MLP head; Acc: accuracy on the pancreatic data. scGL: scGeneLens; Ori: original model; Def: default fine-tuning setting.

| MODEL | VARIANT | EL | ENCODER | MLP | ACC |
| --- | --- | --- | --- | --- | --- |
| SCFOUNDATION | SCGL | – | $L-2$ | ✓ | 0.9625 |
| SCGPT | SCGL | – | ALL | ✓ | 0.8009 |
| SCFOUNDATION | ORI (DEF) | – | $L-2$ | ✓ | 0.9696 |
| SCGPT | ORI (DEF) | ✓ | ALL | ✓ | 0.9625 |
| SCGPT | ORI | – | ALL | ✓ | 0.9133 |
| SCGPT | ORI | – | $L-2$ | ✓ | 0.6698 |

Across scGeneLens hyperparameter settings, scFoundation maintains consistently high cell-type classification accuracy with minimal sensitivity to attention sparsification (Table 1 and Appendix Table 3). In contrast, scGPT is substantially more fragile: its accuracy degrades under scGeneLens and becomes highly sensitive to the number of trainable parameters. For the original scGPT, freezing the embedding layer reduces accuracy, and restricting updates to only the second-to-last encoder layer and MLP head leads to a further drop; similarly, under scGeneLens, performance deteriorates unless more layers are trainable. Consistent with these trends, scFoundation typically converges within ∼ 10 epochs, whereas scGPT often requires more than 25 epochs.

These fine-tuning behaviors suggest that scFoundation and scGPT differ in how explicitly they encode cell identity prior to supervision. To test this, we visualize cell embeddings obtained after MLM fine-tuning only, without supervision by cell-type labels (Fig. 2). scFoundation already separates major cell types into distinct clusters, indicating that strong cell-type structure emerges during MLM and is preserved under scGeneLens-Attention. In contrast, scGPT exhibits much weaker separation, with related endocrine types (e.g., alpha and beta) and lineage-related populations (e.g., ductal and acinar) remaining closer, consistent with its lower annotation performance. We perform the same analysis on a peripheral blood mononuclear cell (PBMC) dataset (Zheng et al., 2017) and find similar patterns (Appendix Fig. 10), suggesting that scFoundation’s stronger intrinsic encoding of cell identity is not dataset-specific.

**Figure 2:**
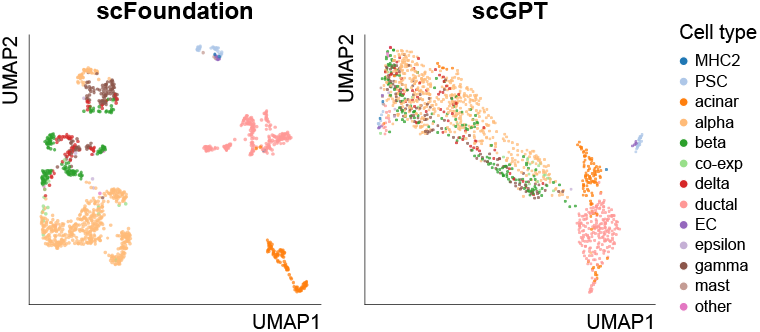
UMAP visualization of pancreas cell embeddings after MLM fine-tuning, without access to cell-type labels.

#### 4.1.2. Retained Activity Information IN SCGPT

To probe which biological signals are preferentially preserved by scGPT under scGeneLens, we analyze a setting in which cell-type identity is controlled. Specifically, we use a fibroblast-only dataset spanning multiple disease contexts (Buechler et al., 2021), including COVID-19, idiopathic pulmonary fibrosis (IPF), non-small cell lung cancer (NSCLC), pancreatic ductal adenocarcinoma (PDAC), and ulcerative colitis (UC), enabling assessment of whether model representations emphasize disease-specific variation or instead capture disease-invariant biological programs.

As shown in Fig. 3, using standard dimensionality reduction, cells in this dataset primarily organize by disease of origin. After MLM fine-tuning with scGeneLens-Attention, scFoundation produces embeddings that clearly separate fibroblasts by disease, indicating a dominant capture of inter-disease variation within a single cell type. In contrast, scGPT does not organize cells by disease; instead, its embedding space forms two major clusters containing cells from all disease contexts, suggesting a latent axis that cuts across disease boundaries.

**Figure 3:**
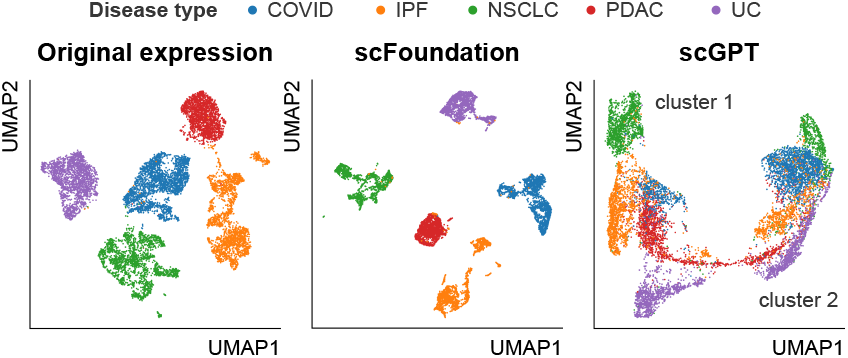
UMAP visualization of original expression and scFM’s embeddings on the fibroblast dataset.

To characterize the functional differences between the two scGPT-derived clusters, we performed Hallmark gene set enrichment analysis (Appendix A.3.1). As shown in Fig. 4, the two clusters display clearly distinct functional profiles. Cluster 1 is enriched for inflammatory and stress-response pathways, including TNF-alpha/NFkB, IL-2/STAT5, interferon gamma signaling, hypoxia, and p53 pathways, suggesting a signal-responsive fibroblast state that primarily reacts to cytokine and environmental cues. In contrast, Cluster 2 shows enrichment of angiogenesis, TGF-beta signaling, and metabolic programs, suggesting a matrix-remodeling, contractile fibroblast population involved in tissue remodeling and structural support. These results indicate that the scGPT-based clustering reflects meaningful biological programs rather than technical artifacts.

**Figure 4:**
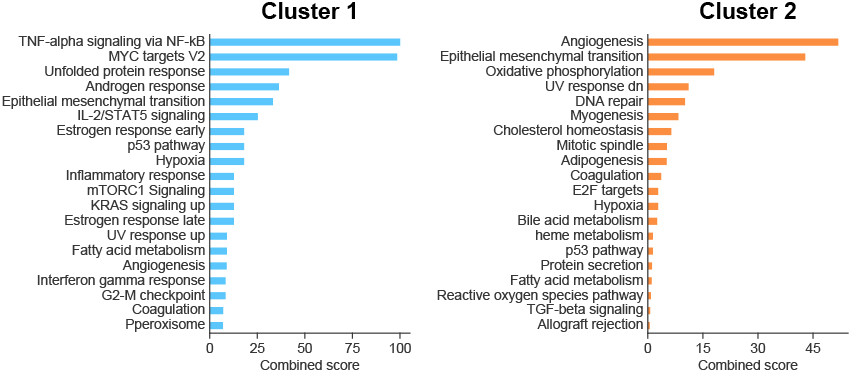
Top Hallmark gene sets enriched among differentially expressed genes between the two clusters in scGPT’s embedding, ranked by combined scores derived by Enrichr.

We next examined whether this functional split is consistent across disease contexts. For each disease subset, we quantified per-cell Hallmark activity differences between the two scGPT-derived clusters (Appendix A.3.2). The results (Fig. 5) reveal strong cross-disease consistency: across COVID-19, IPF, NSCLC, and UC, Cluster 1 consistently exhibits higher activation of stress- and signal-responsive programs, accompanied by elevated MYC- and mTORC1-associated activity. In PDAC, the separation is weaker, consistent with prior reports that cancer-associated fibroblasts are highly plastic and context-dependent with interconvertible states (Biffi et al., 2019). This pattern indicates that scGPT captures a disease-invariant activation axis rather than primarily separating cells by disease of origin, consistent with prior pan-tissue fibroblast atlases (Buechler et al., 2021), which proposed that fibroblast heterogeneity is organized around conserved activation states rather than tissue or disease identity.

**Figure 5:**
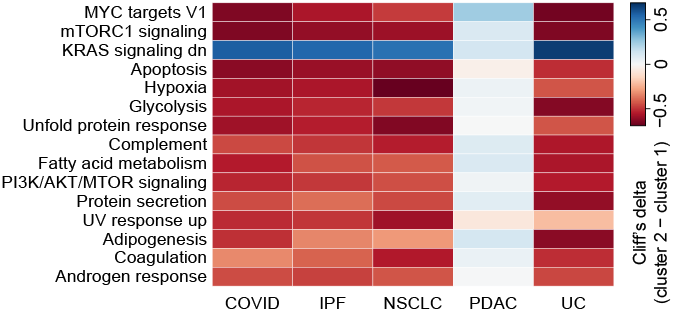
Difference between per-cell Hallmark activity scores and binary cluster membership in scGPT’s embedding, computed separately within each disease subset.

Overall, these results suggest that scGPT preferentially encodes pathway-level activity programs that generalize across conditions, in contrast to scFoundation’s stronger emphasis on condition- or identity-linked variation.

### 4.2. Distinct Gene-level Priors Encoded in Static Token Embeddings

In both scGPT and scFoundation, the Transformer encoder takes as input the sum of a static gene token embedding *E*_*g*_ and an expression value embedding *E*_*e*_. We focus on the static gene embeddings *E*_*g*_, which are learned during pretraining and remain frozen during downstream finetuning, such that differences in *E*_*g*_ directly reflect distinct pretraining-induced inductive biases.

To assess the global organization of static gene embeddings, we visualize the embedding spaces using t-SNE (Fig. 6). scGPT exhibits a markedly more structured embedding space with coherent global organization, whereas scFoundation shows a substantially less organized geometry, consistent with a largely unstructured static embedding space.

**Figure 6:**
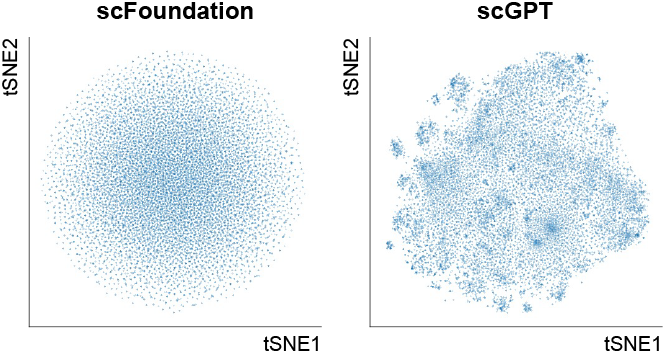
t-SNE visualization of static gene embeddings.

Consistent with this difference, static gene embeddings from scGPT support meaningful functional organization, while those from scFoundation do not (Appendix A.4). Together, these results indicate that scGPT enters the Transformer encoder with a substantially richer gene-level prior encoded in *E*_*g*_, whereas scFoundation’s static token embedding functions only as a gene identity marker.

### 4.3. Tracing Gene-level Information Flow via Influence Propagation

We apply gene influence propagation to trace how genelevel information is integrated across Transformer layers under two settings: MLM fine-tuning only and additional cell-type supervision. At each depth *k*, we construct a propagated influence graph from layer 0 to layer *k*, summarizing cumulative gene–gene influences after stacking the first *k* encoder layers.

#### 4.3.1. Progressive Concentration OF Attention ON A Small Gene Subset

We examine how gene-level attention is redistributed across successive Transformer layers using the propagated influence graph up to depth *k*, denoted as **A**^(*k*)^ ∈ ℝ^*n*×*n*^. For each gene, we quantify incoming propagated influence by the column sum

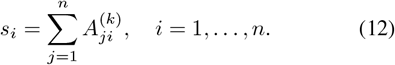

which reflects the extent to which a gene acts as an information sink after stacking the first *k* layers. To assess how unevenly influence is distributed across genes, we compute the Gini coefficient

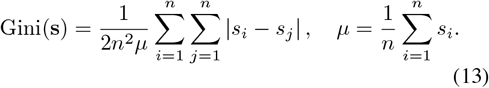

where larger values indicate a stronger concentration of attention onto fewer genes. We additionally report the fraction of total influence captured by the top 0.3% of genes (approximately 20 genes), ranked by incoming propagation, as an interpretable measure of extreme concentration.

Here we take a pancreatic alpha cell as an example. We found that attention concentration increases with depth in both models: the Gini coefficient rises rapidly in early layers and plateaus near 1, indicating that propagated influence is eventually absorbed by a small subset of genes (Fig. 7A). The Top 0.3% metric, however, reveals clear model-specific differences (Fig. 7B). scFoundation reaches high concentration in middle layers and shows a slight reduction toward later layers, whereas scGPT remains less concentrated and increases more gradually. After cell-type supervised finetuning, scGPT primarily exhibits increased concentration in early layers, with only modest changes in deeper layers. In contrast, scFoundation shows only minor early-layer changes, attributable to the introduction of additional special tokens during fine-tuning despite frozen early encoder layers. Similar results are shown in analyzing cells with other types (Appendix Fig. 11A–B).

**Figure 7:**
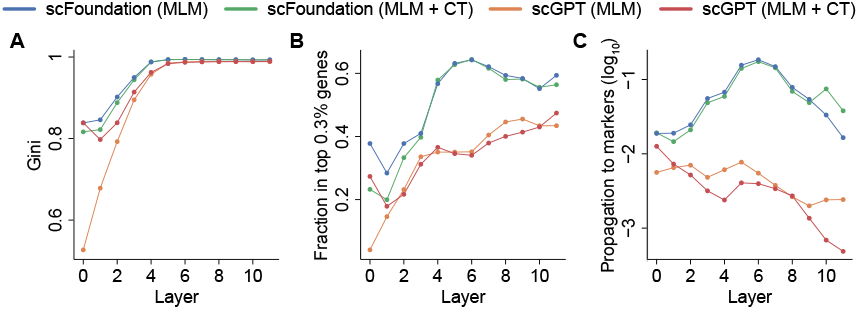
(A–B) Layer-wise concentration of gene influence propagation. (C) Attention propagation to cell marker genes. The results of a pancreatic alpha cell are shown. MLM: fine-tuned by MLM only; MLM + CT: fine-tuned by MLM and cell-type supervision.

#### 4.3.2. Preferential Propagation TOWARD Cell-TYPE Marker Genes

Given the progressive concentration of attention, we next examine which genes act as dominant sinks of propagated influence, focusing on whether cell-type marker genes are preferentially emphasized (details in Appendix Table 6).

As shown in Fig. 7C and Appendix Fig. 11C, scFoundation exhibits strong preferential propagation toward cell-type marker genes. Marker-related influence is weak in early layers, peaks in middle layers, and decreases toward later layers, indicating that marker signals are prominently utilized during MLM fine-tuning, with mid-layer representations playing a central role. This pattern remains largely unchanged after cell-type supervised fine-tuning, with only a modest increase near the second-to-last layer, consistent with scFoundation’s fine-tuning strategy.

In contrast, scGPT shows consistently weaker propagation to marker genes across all layers. Moreover, supervised fine-tuning does not enhance marker-related propagation and can even further reduce it in deeper layers. Together with scGPT’s greater sensitivity to parameter freezing and longer convergence during fine-tuning, these results suggest that scGPT’s MLM representations encode weaker direct cell-type signals, requiring broader representational adjustments rather than amplification of existing marker-driven pathways.

#### 4.3.3. Gene-LEVEL Interpretation OF Influence Propagation Dynamics

To make gene influence propagation patterns biologically interpretable, we examine which genes act as dominant information sinks at different depths after MLM fine-tuning. Specifically, we track the top 1% genes by propagated influence at each layer, illustrating the evolution of gene-level focus using a representative pancreatic alpha cell (the results of a representative acinar cell shown in Appendix A.5).

In scFoundation, propagation centers (genes with the highest propagated attention at each layer) exhibit a clear and orderly transition from cell identity toward functional execution. Early layers emphasize endocrine and secretory signals, with the alpha-cell marker *GCG* and secretory-granule genes (*SCG5, SCG2, CHGB, SCGN*) among the top-ranked genes. In middle layers, accumulation expands toward secretion-related machinery, including transporters such as *ABCC8* and *SLC30A8*, while *GCG* remains prominent. In later layers, attention progressively shifts toward housekeeping and protein homeostasis programs dominated by ribosomal, chaperone, and proteasomal genes, although alpha-cell marker *GCG* can still be observed among the top genes.

In contrast, scGPT exhibits a fundamentally different organization of propagation centers. Early layers are dominated by broad state-related signals rather than endocrine identity, including stress- and immune-associated genes from the *S100* family, *HLA* genes, and inflammatory chemokines. Middle layers emphasize lipid transport and metabolic programs, particularly apolipoproteins (*APOC3, APOA2, APOC1, APOE*). In later layers, propagation becomes highly recurrent, with a small set of genes repeatedly acting as strong sinks, including members of the *ZSCAN, ZSWIM*, and *ZNF* families, together with global response and transport programs.

Overall, the two models organize influence propagation around distinct gene-level centers. scFoundation converges on coherent endocrine modules anchored by canonical marker genes and secretion machinery, yielding an interpretable progression from identity to function. In contrast, scGPT preferentially amplifies family- or state-level gene groups that recur across layers, reflecting coordinated activity programs rather than stable cell-type anchors.

### 4.4. Differences in the Reliance on Gene Identity and Expression Magnitude

To explain the distinct biological signals emphasized by scFoundation and scGPT, we examine which components of the input representation primarily drive their final cell embeddings. Each encoder input consists of a static gene identity embedding *E*_*g*_ and an expression value embedding *E*_*e*_. We apply integrated gradients (IG) to attribute the final cell representation to these two components, aggregating attributions across tokens and embedding dimensions.

As shown in Fig. 8, scGPT consistently assigns a larger fraction of attribution to expression values (*r*_*e*_) than scFoundation, indicating a stronger reliance on *E*_*e*_ in shaping its final representations. In contrast, scFoundation places relatively more attribution on gene identity embeddings *E*_*g*_.

**Figure 8:**
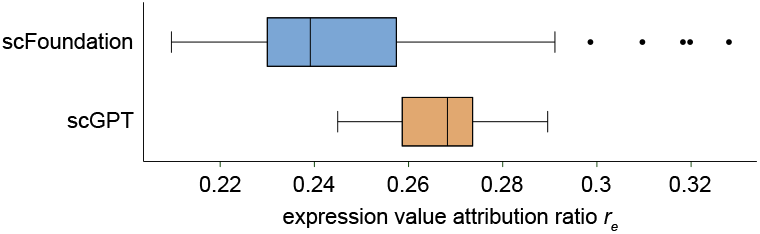
Comparison of expression value contribution ratio (*r*_*e*_) between scGPT and scFoundation. For each cell in the pancreatic dataset, we compute the fraction of attribution mass assigned to the expression value embedding.

This attribution difference provides a coherent explanation for the contrasting behaviors observed throughout our analyses. Expression values more directly reflect pathway activity and cell states, consistent with scGPT’s tendency to organize representations around disease-invariant activation programs and broad state signals. Conversely, greater reliance on gene identity embeddings aligns with scFoundation’s emphasis on marker genes and its progressive propagation from cell identity toward functionally coherent modules.

Finally, this attribution gap likely reflects deeper differences established during pretraining. scGPT’s static gene token embeddings already encode substantial biological structure, whereas scFoundation’s static embeddings appear comparatively unstructured, placing a greater burden on attention-mediated integration. How pretraining objectives, data composition, and modeling choices jointly shape the balance between gene identity and expression value signals remains an open question for future work.

## 5. Discussion and Conclusions

In this work, we introduced scGeneLens, an attention-based framework for dissecting how scFMs encode biological information. By systematically comparing scFoundation and scGPT, we show that scFMs with similar Transformer architectures can rely on fundamentally different biological signals. scFoundation organizes representations around cell identity and context-defining marker programs, whereas scGPT emphasizes pathway-level activity patterns that generalize across conditions. These findings demonstrate that interpretability is essential for revealing the inductive biases underlying scFMs.

Our analyses further indicate that architectural similarity does not imply similarity in the biological abstractions learned by scFMs. Instead, different models adopt distinct organizing principles for cellular representations. scFoundation builds stable, gene-identity–anchored structures in which canonical marker genes and coherent functional modules act as persistent anchors throughout attention propagation. In contrast, scGPT organizes representations around dynamic, pathway-level activity programs that are largely invariant to cell type or disease context. These behaviors correspond to two complementary notions of cell state: a discrete identity defined by marker programs and a continuous position along latent activity axes.

Viewed through this lens, the observed differences should not be interpreted along a single notion of model quality, but rather as trade-offs in how biological information is abstracted and prioritized. Models that emphasize gene identity are better suited for tasks centered on cell-type definition and annotation, whereas models that rely more heavily on expression magnitude naturally capture functional state variation and cross-condition generalization. This perspective helps explain the heterogeneous behavior of scFMs across tasks and suggests that model selection should be guided by the biological question of interest.

### Methodological implications

An important methodological consideration in our approach is the use of block sparsity to facilitate interpretability. When attention is analyzed directly on raw inputs, attribution patterns are highly diffuse, with very low Gini coefficients (Appendix Fig. 12), indicating that attention mass is broadly distributed across many functionally related genes and thus obscures interpretable structure. Introducing block sparsity substantially increases attribution concentration by compressing broadly distributed attention over functionally related genes into a small set of representative genes. This concentration enables clearer feature attribution and more interpretable biological programs, making coherent biological structure more readily discernible and underscoring the necessity of this design.

### Limitations

Our study has several limitations. First, although scGeneLens is broadly applicable to Transformer-based scFMs, analyzing Performer-style models (e.g., scBERT) requires converting approximate attention to standard attention, substantially increasing memory and computational cost. Besides, for models whose input ordering depends on expression values (e.g., Geneformer), the current block-partition strategy may be poorly aligned with the input structure, limiting the suitability of scGeneLens and weakening the reliability of derived conclusions. In addition, scGeneLens currently orders genes lexicographically before block partitioning. While this simple, model-agnostic strategy can partially reflect gene families or shared nomenclature, it does not ensure functional or regulatory coherence within blocks, motivating future exploration of more biologically informed grouping schemes. Finally, biological interpretation is inherently constrained by current domain knowledge, and some gene-level attributions may be incomplete or subject to misinterpretation.

### Future directions

An important next step is to understand how architectural and pretraining choices in scFMs give rise to the distinct inductive biases revealed here, and how these biases shape performance across downstream tasks. More broadly, our results highlight the need for closedloop frameworks that integrate model training, interpretability, task-specific evaluation, and iterative refinement. Such feedback-driven approaches will be critical for moving beyond one-size-fits-all foundation models toward principled, purpose-aligned design of single-cell models.

## Impact Statement

This paper aims to improve the understanding and interpretability of single-cell foundation models. While the proposed framework may facilitate biological discovery and downstream biomedical research, we do not foresee any immediate ethical concerns beyond those generally associated with machine learning research.

## A. Appendix

### A.1. scGeneLens Setup and Hyperparameters

We apply scGeneLens-Attention as a drop-in replacement for standard self-attention in the encoder, controlled by two hyperparameters: TopK and chunk size. TopK specifies the number of blocks retained for each query at the block-selection stage, while chunk size determines the number of gene tokens grouped into each block. TopK directly controls the sparsity of the attention pattern: smaller values retain fewer high-attention connections, improving interpretability but potentially reducing model performance. Chunk size affects both computational cost and the degree of stochastic connectivity introduced by block-wise partitioning. In practice, very small chunk sizes may consistently block a subset of genes, whereas excessively large chunk sizes introduce too many weak connections and reduce the clarity of attention structure.

Based on these considerations, we conduct a grid search over chunk sizes {16, 32, 64, 128} and TopK values {4, 6, 8, 12, 24}. For each hyperparameter configuration, we fine-tune the model using the masked language modeling (MLM) objective and report the resulting MLM loss in Appendix Table 2. The cell-type annotation accuracies of different scFMs under varying TopK and chunk sizes on the pancreatic dataset are summarized in Appendix Table 3.

Based on the downstream cell-type annotation results, we found that TopK = 4 with chunk size = 16 led to a clear performance degradation compared with TopK = 6 with chunk size = 16. Moreover, the performance of TopK = 6 with chunk size = 16 was comparable to that obtained with larger TopK settings, while also yielding a model that is easier to interpret. Therefore, we adopt the model with TopK = 6 and chunk size = 16 as our default setting for model interpretation in all subsequent experiments.

**Table 2:**
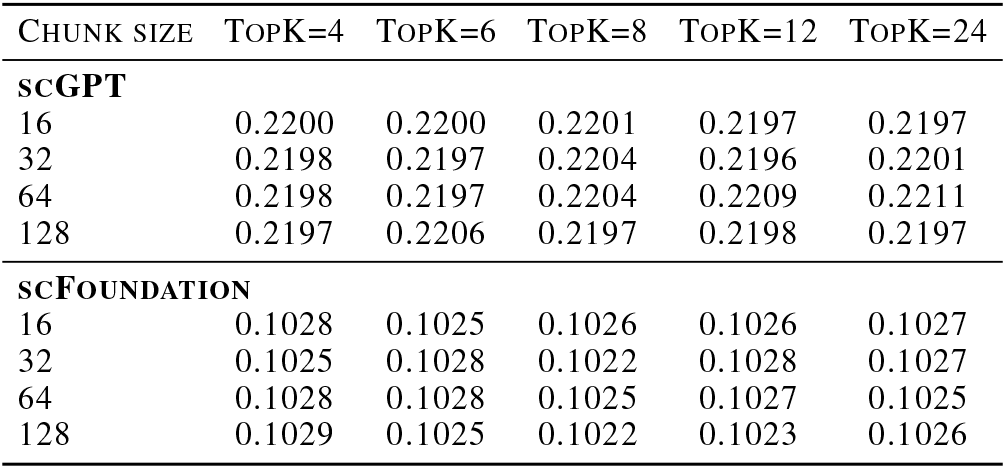
MLM loss of scGPT and scFoundation fine-tuned with scGeneLens under different TopK and chunk-size settings. Lower values indicate better performance.

### A.2 Datasets

We evaluate scGeneLens on multiple single-cell RNA-seq datasets covering diverse biological contexts. For cell-type annotation analyses, we use a human pancreatic dataset (Segerstolpe et al., 2016) and a peripheral blood mononuclear cell (PBMC) dataset (Zheng et al., 2017). The pancreatic dataset contains 13 annotated cell types, including major pancreatic populations such as alpha, beta, delta, gamma/PP, acinar, and ductal cells. The PBMC dataset comprises 11 annotated immune cell types.

To further examine model behavior across tissues and disease conditions within a single cell type, we additionally analyze a human fibroblast-only dataset (Buechler et al., 2021). This dataset includes fibroblasts collected from multiple pathological contexts, including COVID-19, idiopathic pulmonary fibrosis (IPF), non-small cell lung cancer (NSCLC), pancreatic ductal adenocarcinoma (PDAC), and ulcerative colitis (UC).

### A.3. Interpreting information encoded in scFMs

#### A.3.1. Hallmark Gene Set Enrichment

We identify differentially expressed genes between the two fibroblast clusters, perform Hallmark gene set enrichment using Enrichr (Kuleshov et al., 2016) with the MSigDB Hallmark collection (Liberzon et al., 2015). Genes are stratified by the direction of differential expression, and enrichment results are ranked by the Enrichr combined score.

**Table 3:** Cell type annotation accuracy of scGPT and scFoundation fine-tuned with scGeneLens across different TopK and chunk size settings on the pancreatic dataset.

| CHUNK SIZE | TOPK=4 | TOPK=6 | TOPK=8 | TOPK=12 | TOPK=24 |
| --- | --- | --- | --- | --- | --- |
| **scGPT** |  |  |  |  |  |
| 16 | 0.6159 | 0.8009 | 0.7541 | 0.7658 | 0.8548 |
| 32 | 0.7377 | 0.6979 | 0.7377 | 0.7752 | 0.8173 |
| 64 | 0.7096 | 0.8080 | 0.7330 | 0.7541 | 0.8384 |
| 128 | 0.8126 | 0.8337 | 0.8454 | 0.8314 | 0.7705 |
| **SCFOUNDATION** |  |  |  |  |  |
| 16 | 0.9555 | 0.9649 | 0.9602 | 0.9555 | 0.9578 |
| 32 | 0.9532 | 0.9555 | 0.9649 | 0.9555 | 0.9602 |
| 64 | 0.9555 | 0.9672 | 0.9649 | 0.9555 | 0.9532 |
| 128 | 0.9672 | 0.9649 | 0.9532 | 0.9649 | 0.9578 |

#### A.3.2. Quantifying Hallmark Activity Differences Between SCGPT-Derived Clusters

We quantify differences in Hallmark activity between scGPT-derived clusters using Cliff’s delta. Single-cell Hallmark scores are computed by aggregating expression over genes in each Hallmark set, and Cliff’s delta is calculated between cluster-specific score distributions within each disease context, with the sign indicating direction and the magnitude reflecting effect size.

### A.4. Functional Organization of Static Gene Embeddings

To assess whether static gene embeddings encode gene-level functional organization, we formulated a binary classification task based on Gene Ontology (GO) annotations. For each selected GO term, we constructed positive gene pairs from genes annotated to the same term and sampled an equal number of negative gene pairs from genes not sharing that annotation. To control for sampling bias, the same set of negative pairs was used across different embedding models within each experiment.

For each GO term, the resulting dataset was randomly split into 80% for training and 20% for testing. This procedure was repeated ten times with different random seeds to assess robustness. Static gene embeddings were standardized using z-score normalization prior to classification. We trained a linear support vector machine (SVM) classifier on top of the static gene embeddings to predict whether a given gene pair belonged to the same GO term. Model performance was evaluated using the area under the receiver operating characteristic curve (AUC), and the distribution of AUC values across repeated runs was summarized using boxplots.

As shown in Appendix Fig. 9, classifiers trained on scGPT static gene embeddings consistently achieve higher AUC values across a wide range of GO terms. This indicates that functional relationships defined by GO annotations are partially separable in the scGPT embedding space. In contrast, classifiers trained on scFoundation static gene embeddings perform close to chance level (AUC ≈ 0.5) for most GO terms, suggesting that functional similarity between genes is not explicitly reflected in its static embedding geometry.

**Figure 9:**
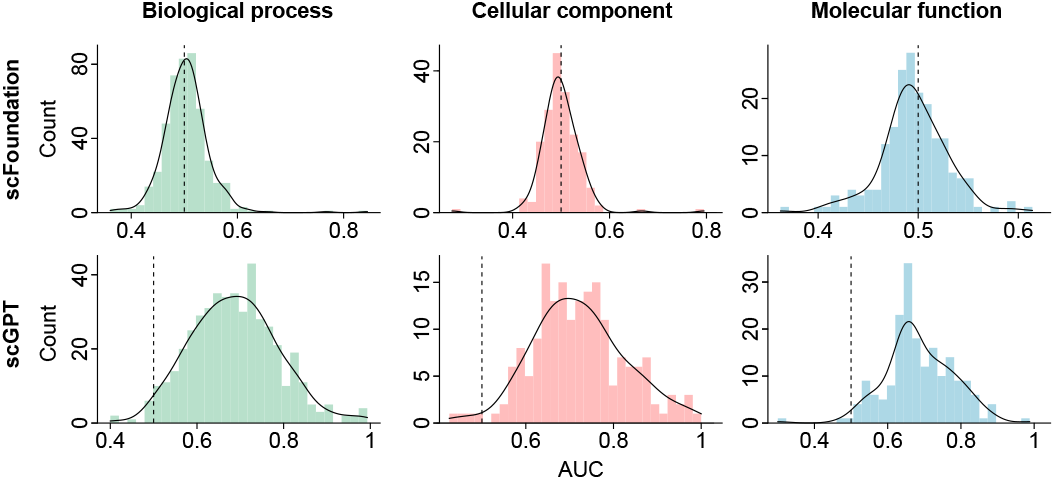
SVM classification performance (AUC) for predicting GO term memberships using static gene embeddings from different scFMs. Dashed lines denote chance-level performance (AUC = 0.5).

Notably, this evaluation relies solely on static gene token embeddings, without incorporating contextualized representations or cell-level information. The observed performance differences therefore reflect intrinsic properties of the pre-trained embedding spaces, rather than effects introduced by downstream supervision or task-specific fine-tuning.

### A.5. Gene-level Interpretation of A Pancreatic Acinar Cell

#### scFoundation

In scFoundation, the propagation centers for the acinar cell reveal a structured depth-wise progression from strong cell-identity anchoring to increasingly general cellular programs. We get the following analysis from Appendix Table 4.

In the early layers, propagated attention is dominated by canonical acinar markers and secretion-associated genes, indicating that the model routes influence toward cell-type–defining signals from the start. Prominent centers include digestive enzyme genes such as *CTRB1* and *CTRB2*, together with regeneration- and acinar-enriched genes from the *REG* family, including *REG1A, REG3A*, and *REG1B*. These identity anchors co-occur with epithelial and secretion-adjacent signals such as *MUC1* and *MUC20*, as well as protease-inhibitor and stress-related genes such as *SERPINA3*. In parallel, the early layers also highlight components consistent with high secretory burden, including trafficking and membrane factors and early signs of ER–Golgi organization, suggesting that shallow representations already integrate acinar identity with the infrastructure required for protein production and secretion.

In the middle layers, the acinar identity anchors remain among the strongest propagation centers, while influence increasingly spreads into machinery that supports secretion and protein handling. Trafficking and secretory-pathway components become more visible, including ER–Golgi and vesicle-transport genes from the *TMED* and *SEC* families and related transport factors. At the same time, protein homeostasis modules gain prominence, with more ribosomal genes, chaperones, and lysosomal proteases such as *CTSB* and *CTSD* appearing among the propagation centers. In addition, state-related signals including chemokines such as *CXCL12, CXCL6*, and *CXCL14* can enter the top lists, consistent with the model capturing not only identity but also microenvironmental or stress-associated variation in acinar states.

In the later layers, the propagation centers shift more strongly toward housekeeping and metabolic programs, reflecting a move to broad cellular maintenance features. The top centers become enriched for ribosomal proteins, mitochondrial components, and RNA or protein processing factors, together with continued emphasis on proteostasis-related genes.

#### scGPT

In scGPT, the propagation centers are dominated by coordinated state programs rather than acinar-specific secretion modules, and they evolve toward highly recurrent sink-like gene families in deeper layers. We get the following analysis from Appendix Table 5.

In the early layers, propagation concentrates on immune and inflammatory signals. The top centers are enriched for chemokines such as *CXCL2, CXCL3*, and *CXCL6*, together with antigen-presentation genes from the *HLA* locus including *HLA-DRA, HLA-DRB1, HLA-DQB1*, and *HLA-DMB*. Stress-associated families also appear prominently, including *S100A13, S100A14*, and multiple metallothioneins such as *MT1M, MT1E, MT1X*, and *MT2A*. By layers 2–3, additional state axes enter the propagation centers, including endothelial and signaling components such as *PLVAP, NOS3*, and *NOTCH4*, alongside extracellular matrix genes such as *COL6A1* and *COL6A2*, indicating a shift from purely inflammatory programs toward a mixed immune–endothelial–matrix state description.

In the middle layers, propagation increasingly converges on a stable set of recurrent sink-like families, led by *ZSCAN31, ZSCAN26, ZSCAN5B, ZSWIM1, ZSWIM5, ZWINT, ZW10*, and *ZZEF1*, together with *ZYX*. These centers co-occur with persistent immune markers from the *HLA* locus, chemokines such as *CXCL1, CXCL2, CXCL12*, and *CXCL16*, and stress-related metallothioneins. At the same time, matrix-associated genes become increasingly prominent, including *COL2A1, COL4A1*, and *COL18A1*, suggesting that intermediate-depth representations emphasize structured convergence and extracellular matrix organization.

In the later layers, extracellular matrix signatures become a primary axis of propagation centers, with broad enrichment of collagen genes including *COL1A1, COL1A2, COL3A1, COL11A2, COL12A1, COL13A1, COL14A1, COL15A1, COL16A1, COL18A1, COL21A1, COL23A1*, and *COL27A1*. In parallel, glycosaminoglycan and matrix-modifying enzymes such as *CHSY3* and sulfotransferases including *CHST1, CHST8, CHST13, CHST3*, and *CHST4* appear among the top centers, while the recurrent sink families *ZSCAN, ZSWIM, ZWINT, ZW10*, and *ZZEF1* remain consistently ranked. Overall, scGPT’s depth-wise propagation centers are organized around co-activated immune, endothelial, and extracellular matrix programs, with later layers characterized by strong structured convergence onto recurrent sink-like gene families.

### A.6. Appendix Tables and Figures

**Figure 10:**
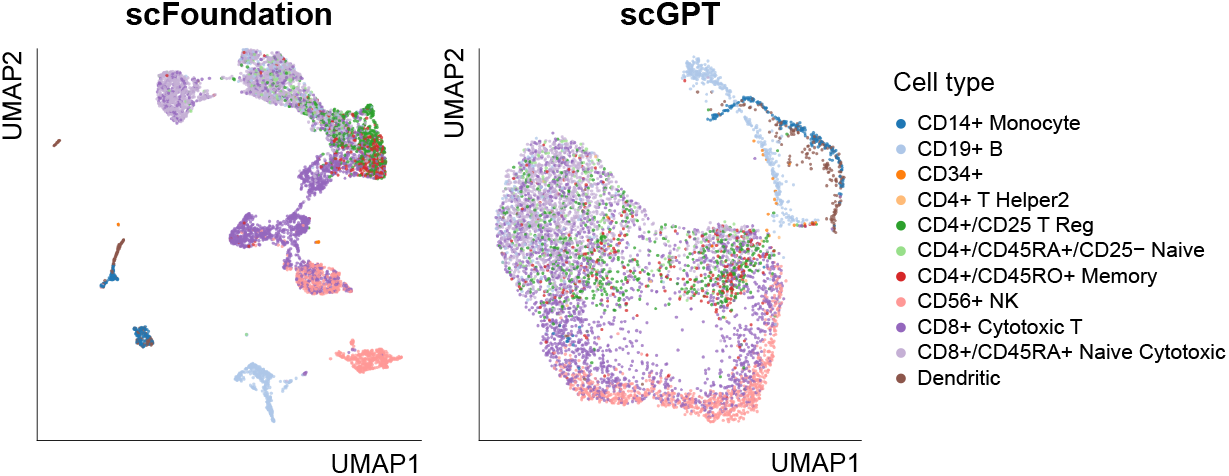
UMAP visualization of PBMC embeddings after MLM fine-tuning, without access to cell-type labels.

**Figure 11:**
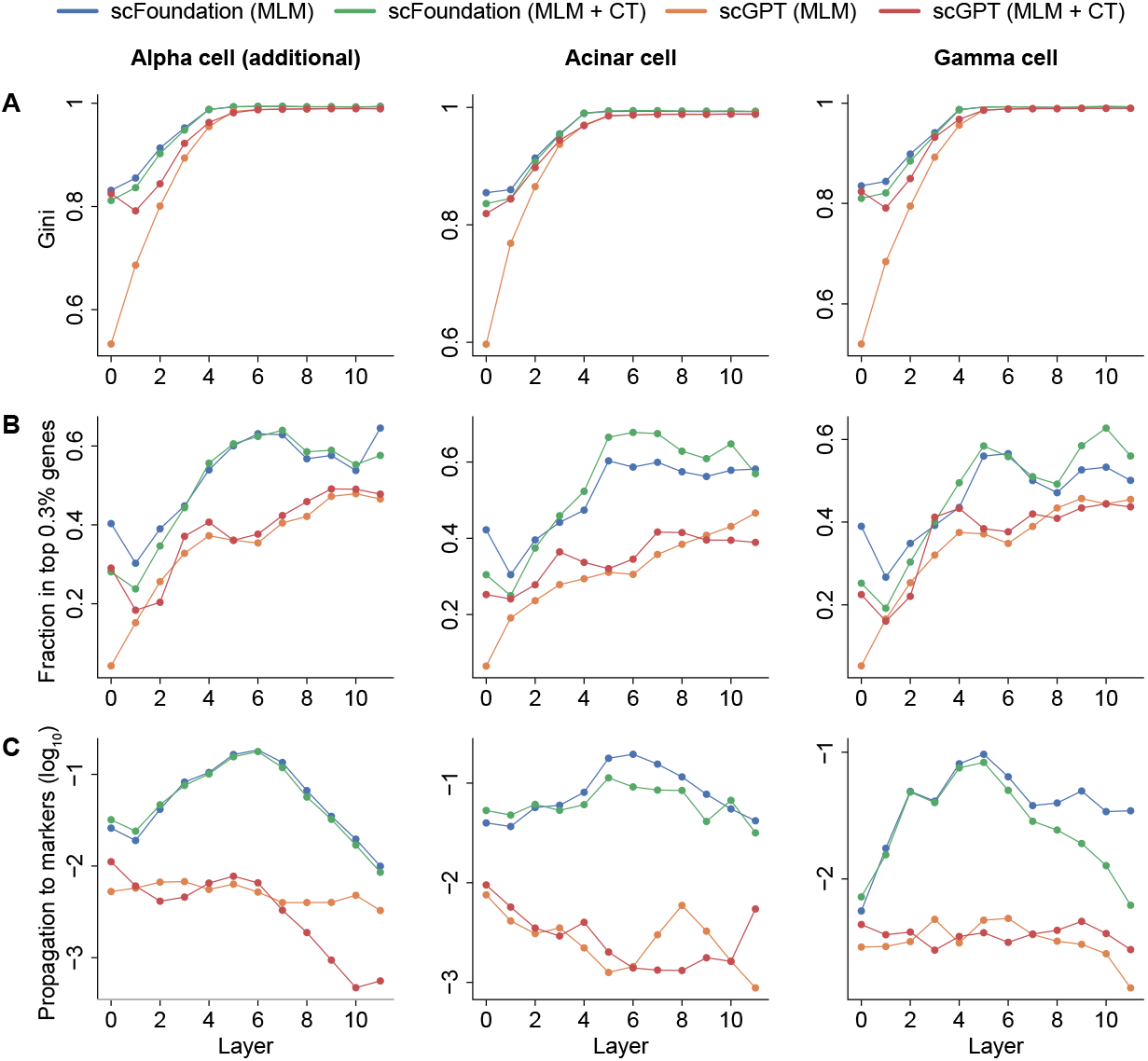
(A–B) Layer-wise concentration of gene influence propagation. (C) Attention propagation to cell marker genes. The results of an additional pancreatic alpha cell, an acinar cell, and a gamma cell are shown. MLM: fine-tuned by MLM only; MLM + CT: fine-tuned by MLM and cell-type supervision.

**Figure 12:**
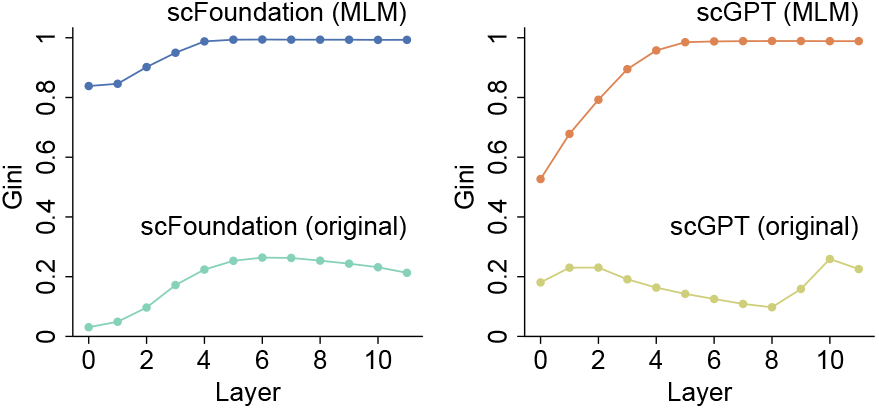
Layer-wise concentration of propagated gene influence in original scFMs and their block-sparse counterparts derived using scGeneLens-Attention under MLM fine-tuning.

**Table 4:**
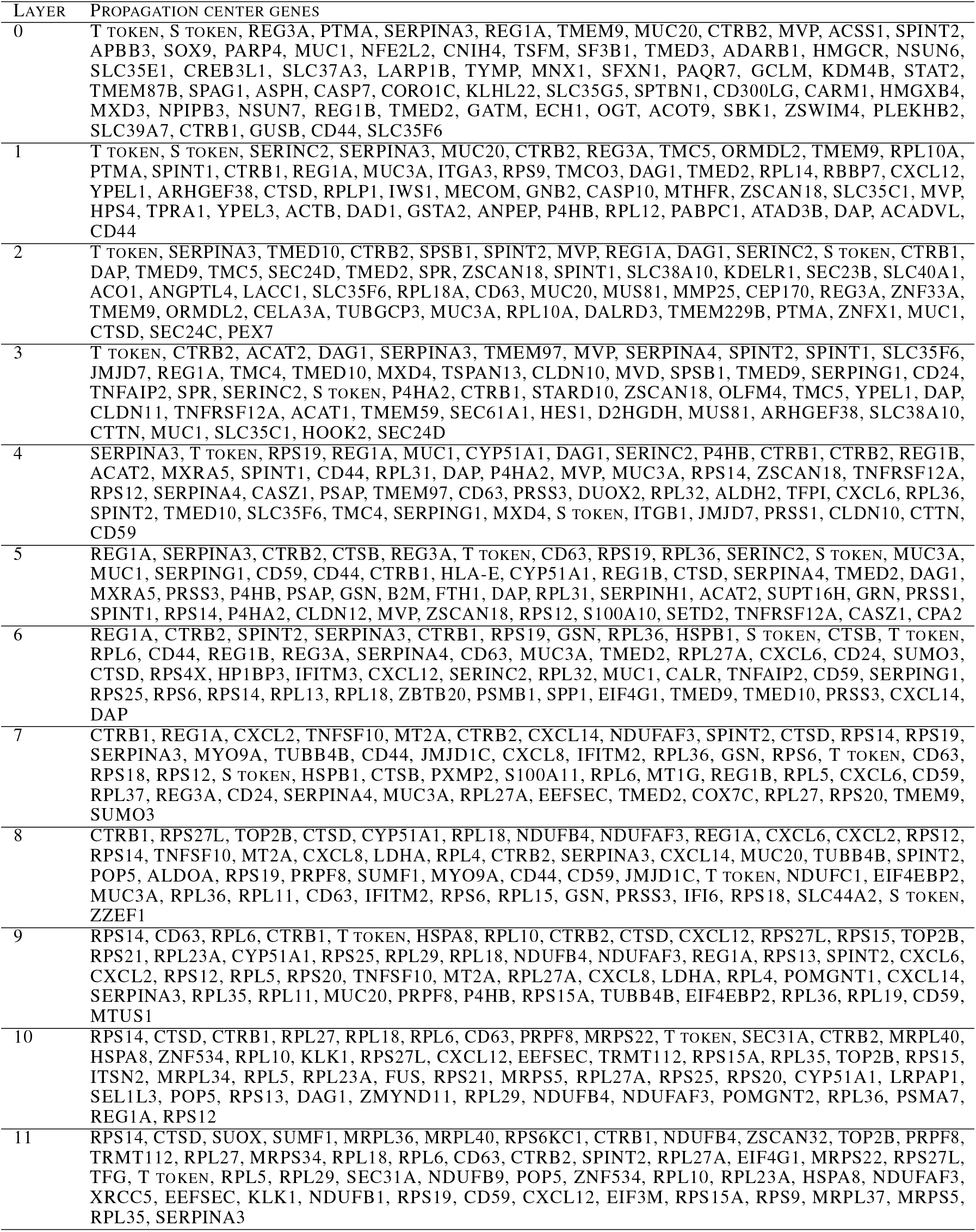
Propagation centers of scFoundation in different layers for a pancreatic acinar cell.

**Table 5:**
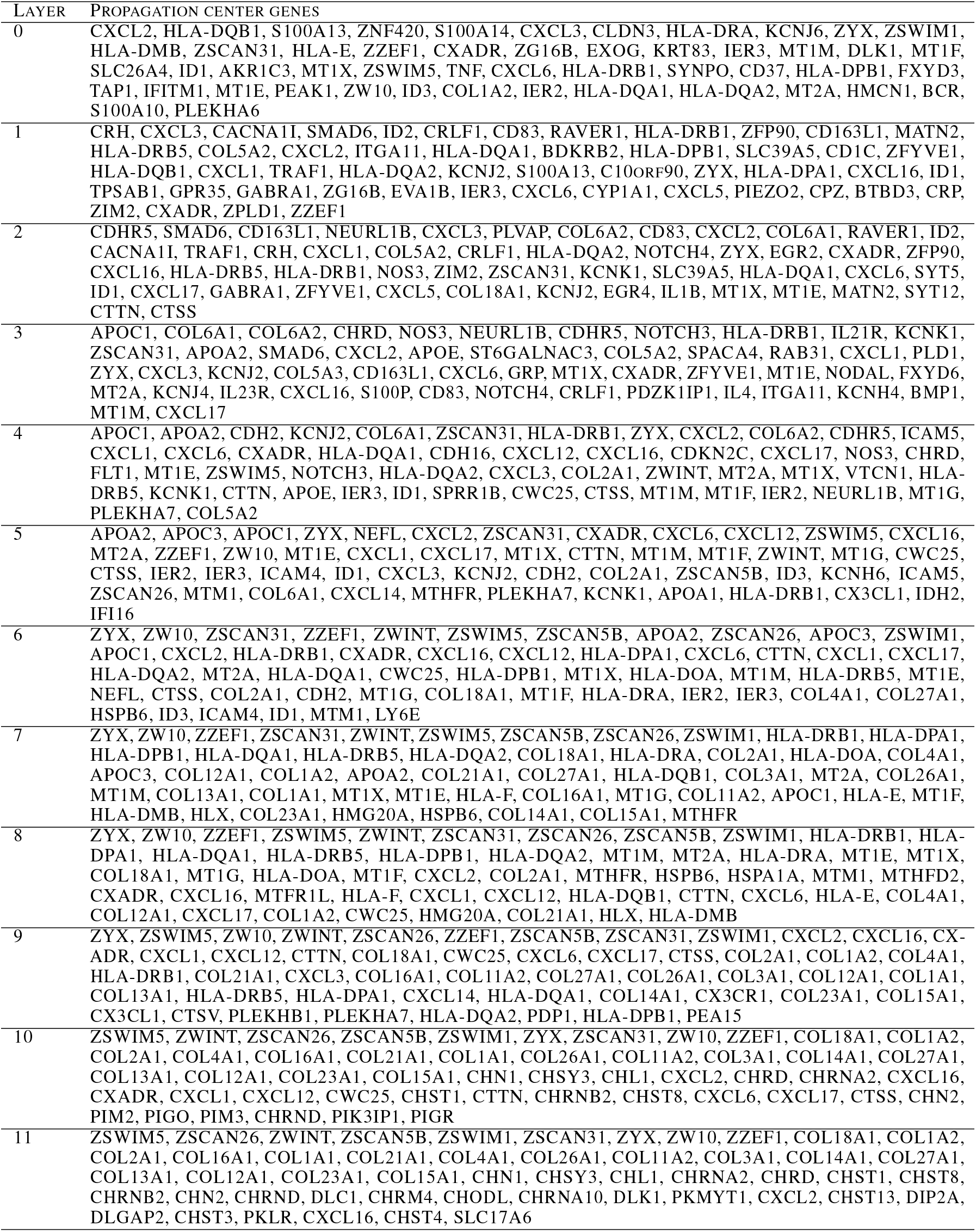
Propagation centers of scGPT in different layers for a pancreatic acinar cell.

**Table 6:** Marker genes of different cell types.

| TYPE | MARKER GENES |
| --- | --- |
| ALPHA CELL | GCG, ARX, TTR, IRX2, MAFB, PAX6, NEUROD1, CHGA, SCGN, RESP18, PCSK1, PCSK2, GC, ALDH1A1, UCP2, RGS4, KCNK16, FEV, RFX6, SLC30A8, SMIM24, PEMT, PRRG2, PLCE1, IRX1, TM4SF4, CRYBA2, LOXL4 |
| ACINAR CELL | PRSS1, PRSS2, PRSS3, CTRB1, CTRB2, CTRC, CPA1, CPA2, CPB1, CELA3A, CELA3B, CELA2A, CELA2B, CELA1, AMY1A, AMY1B, AMY2A, AMY2B, PNLIP, CLPS, CEL, SYCN, REG1A, REG1B, REG3A, SPINK1, RBPJL, PTF1A, BHLHA15, NR5A2, XBP1, GP2, ERP27, FOXP2 |
| GAMMA CELL | PPY, CARTPT, GAD2, PCSK1N, SCG2, SLC6A4, IL17RB, PAX6, ISL1, NEUROD1, NKX2-2, ETV1, FEV, TOX3, CALB1, STMN2, GRIA3, CHRM3, KCNJ3, KCNJ8, KCNMB2, CHGA, CHGB, PCSK1, PCSK2, SYP |

## Notes

### Competing Interest Statement

The authors have declared no competing interest.

### Funding

- The work is sponsored by National Natural Science Foundation of China (92470105, 62373210), the National Key R&D Program of China (2025YFC3409300).

